# Failure of colonization following gut microbiota transfer exacerbates DSS-induced colitis

**DOI:** 10.1101/2024.09.25.614792

**Authors:** Kevin L. Gustafson, Trevor R. Rodriguez, Zachary L. McAdams, Lyndon M. Coghill, Aaron C. Ericsson, Craig L. Franklin

## Abstract

To study the impact of differing specific pathogen-free gut microbiomes (GMs) on a murine model of inflammatory bowel disease, selected GMs were transferred using embryo transfer (ET), cross-fostering (CF), and co-housing (CH). Prior work showed that the GM transfer method and the microbial composition of donor and recipient GMs can influence microbial colonization and disease phenotypes in dextran sodium sulfate-induced colitis. When a low richness GM was transferred to a recipient with a high richness GM via CH, the donor GM failed to successfully colonize, and a more severe disease phenotype resulted when compared to ET or CF, where colonization was successful. By comparing CH and gastric gavage for fecal material transfer, we isolated the microbial component of this effect and determined that differences in disease severity and survival were associated with microbial factors rather than the transfer method itself. Mice receiving a low richness GM via CH and gastric gavage exhibited greater disease severity and higher expression of pro-inflammatory immune mediators compared to those receiving a high richness GM. This study provides valuable insights into the role of GM composition and colonization in disease modulation.

## Introduction

The gut microbiome (GM), the microorganisms that inhabit the gastrointestinal tract of humans and animals, plays an important role in health and disease pathogenesis and severity^1,2^. Common diseases that may be influenced by features within the GM include inflammatory bowel disease (IBD)^3^, colon cancer^4^, and autism^5^ among others. In the case of IBD, changes in the GM characterized by reduced richness of symbiotic commensals (i.e., dysbiosis) may exacerbate inflammation^6^. This dysbiosis induces further immune reactions and inflammation characterized by the production of immune mediators, reactive oxygen species, and antimicrobial peptides when pathobiont microbes breach the mucosa^7,8^. While the GM plays an important role in both physiology and pathophysiology of many diseases including IBD, studying the GM in humans can be cumbersome and difficult due to varying backgrounds of individuals, unknown genetic contribution to disease^9^ and the correlative nature of human studies. To overcome these challenges, animal models have been established to answer questions concerning how the microbiome interacts with and influences the health of the host. Studies performed in gnotobiotic and germ-free rodents have established the need for a healthy GM for proper biological development and physiology of the host organism^10–12^. While animal models are essential to the advancement of scientific knowledge, many studies utilizing rodents as models suffer from poor reproducibility or translatability^13,14^. Awareness of these issues has led the National Institutes of Health to launch an initiative to improve the translatability and reproducibility in research to make more investigators and their laboratories aware of the need to improve scientific rigor^15^.

The composition of the GM has an impact on the disease state of the rodent host^16–19^. The rodents provided to the research community by the largest suppliers are colonized by GMs that significantly differ in alpha and beta-diversity^20,21^. Furthermore, research performed in isogenic mice where the experimental groups harbor these different supplier-origin GMs have shown that the GM can influence disease severity independent of genetic contributions in a chronic dextran-sodium sulfate (DSS)-induced murine model of IBD^22^. Previous work in our lab has shown that when co-housing is used to transfer a complex GM, colonization by the donated GM is less successful and may result in severe DSS-induced disease and mortality. We sought to replicate these findings in an acute model of DSS colitis and confirm that the effect on disease severity is attributable to the GM and not an unexplained factor of the co-housing method. We leveraged the differences in GM alpha and beta diversity between a Jackson Laboratory-origin GM, and Envigo-origin GM^23^. Specifically, we hypothesized that the transfer of donor microbiome to recipients naturally during the post-partum period (i.e., via embryo transfer (ET) of recipient germplasm in surrogate dams harboring the donor GM) or within the first 24 hours of life (i.e., via cross-foster (CF) of recipient pups on surrogate dam donors) would result in more complete colonization of the donor GM than co-housing for one month beginning at weaning.

We also hypothesized that attempts to transfer a comparably low-richness GM to recipients harboring a high-richness GM via CH would result in particularly low transfer efficacy, and the most severe disease when challenged with DSS at seven weeks of age. During the CH process, donor and recipient mice were grouped together at weaning (21 days) in cages containing two donors and two recipients, a procedure that could produce inadvertent effects on recipient mice via psychosocial stressors or other unknown factors. To obviate any potential effects of physical contact between the donor and recipient mice during the CH procedure, we included additional treatments groups with the same timing and donor and recipient GMs, wherein the GM was transferred via weekly gastric gavage and dirty bedding transfer. Recipient mice receiving the reciprocal GM via ET, CF, CH, or gastric gavage and transfer of dirty donor bedding (GA) were challenged with a single cycle of DSS, followed by assessment of disease severity via weight loss and mortality, histological examination, colon length at necropsy, and production of inflammatory mediators. Supporting a primary influence of poorly colonizing microbes rather than factors associated with direct physical contact between donors and recipients, we hypothesized that mice receiving a low-richness GM via GA and CH would have equivalent, severe DSS-induced disease compared to all other groups. Conversely, if mice receiving the low-richness GM via CH developed more severe disease than mice receiving the same GM via GA, it would indicate differences in disease severity and survival are associated at least in part with physical contact between recipients and donors.

## Methods

### Ethics Statement

All activities and experiments described using animal models were performed in accordance with the Guide for the Care and Use of Laboratory Animals, and the Institutional Animal Care and Use Committee of the University of Missouri, an AAALAC-accredited institution, approved all animal use procedures (MU IACUC protocols 9587 and 36781).

### Mice

C57BL/6J (B6J) and C57BL/6NHsD (B6N) mice were procured directly from The Jackson Laboratory (Bar Harbor, ME) or Envigo (now Inotiv, Indianapolis, IN), respectively, and bred to produce GM recipient mice. Embryo transfer, cross-fostering, and co-housing transfer methods were performed as previously described^22^. Colonies of mice were housed under barrier conditions in microisolator cages with compressed pelleted paper bedding and nestlets, on ventilated racks with *ad libitum* access to 5053 (LabDiet, St. Louis, MO) rodent chow and acidified, autoclaved water. Mice were housed under a 12:12 light/dark cycle. Mice were determined to be free of bacterial pathogens including *Bordetella bronchiseptica*, *Filobacterium rodentium*, *Citrobacter rodentium, Clostridium piliforme, Corynebacterium bovis, Corynebacterium kutscheri, Helicobacter* spp., *Mycoplasma* spp., *Rodentibacter* spp., *Pneumocystis carinii, Salmonella* spp., *Streptobacillus moniliformis, Streptococcus pneumoniae*; adventitious viruses including H1, Hantaan, KRV, LCMV, MAD1, MHV, MNV, PVM, RCV/SDAV, REO3, RMV, RPV, RTV, and Sendai viruses; intestinal protozoa including *Spironucleus muris, Giardia muris, Entamoeba muris,* trichomonads, and other intestinal flagellates; intestinal helminths including pinworms and tapeworms; and external parasites including all species of lice and mites, via quarterly sentinel testing.

#### CD-1 donor mice

All CD-1 mice that were used as donors were from two colonies in which the founders were originally purchased from Charles River (Crl:CD1(ICR), Frederick, MD), and were generated via rederivation to harbor either a high richness Envigo origin GM (GM^High^), or a low richness Jackson Laboratory origin GM (GM^Low^) as previously described^24^. All donor mice were reared at the authors’ institution and the two colonies have been maintained and continually monitored for GM stability within our facility for over 35 generations. Additionally, a rotational breeding scheme and routine introduction of CD-1 genetics via embryo transfer from CD-1 mice purchased from Charles River allows for the maintenance of allelic heterozygosity within each colony and ensures these colonies do not become genetically distinct from each other. Since CD-1 mice that harbor a Jackson Laboratory origin GM were found to have a GM with low phylogenetic richness and diversity, the GM of these mice was designated GM^Low^. Similarly, since CD-1 mice that harbored an Envigo origin GM were found to have a GM with high phylogenetic richness and diversity relative to GM^Low^, the GM of these mice was designated GM^High^.

#### Embryo transfer

ET was performed at the authors’ institution as previously described^23^. Briefly, B6J and B6N mice were obtained directly from respective producers, bred, and embryos were collected at the two-cell stage. Embryos were surgically transplanted into GM donor pseudopregnant CD-1 surrogate dams. GM donor surrogate CD-1 mice were allowed to give birth and rear the pups (**Sup Fig S1A,B**). Pups that were generated via ET were then bred together to generate a second generation to simulate the breeding that is sometimes necessary to obtain enough animals to power scientific experiments due to low animal yields from embryo transfer procedures^25^. Numbers obtained from these breedings are as follows: GM^High^ET (*n* = 12/sex), GM^Low^ET (*n* = 12/sex). Both male and female mice were included in the ET groups at a 1:1 ratio.

#### Cross-fostering

B6J and B6N mice, obtained directly from respective producers, were bred to generate GM recipient mice. CD-1 dams were time-mated simultaneously with the B6J and B6N dams to act as cross-foster surrogate dams. B6J and B6N pups were cross-fostered to a CD-1 GM donor dam harboring high richness GM^High^ or low richness GM^Low^, respectively, within 12 hours following birth (**Sup Fig S1C,D**). Numbers obtained from the cross-fostering procedure are as follows: GM^High^CF (*n* = 12 males, 11 females), GM^Low^CF (*n* = 12/sex). To limit the possibility of cannibalism and help facilitate GM transfer, 2-3 CD-1 pups born to the surrogate dams remained within the litters.

#### Co-housing

B6J and B6N mice, obtained directly from respective producers, were bred to generate GM recipient mice. CD-1 dams were time-mated simultaneously with the B6J and B6N dams to generate GM donor mice. At 21 days of age, recipient B6J and B6N mice were weaned and co-housed with weanling CD-1 mice harboring GM^High^ or GM^Low^, respectively (**Sup Fig S1E-H**). B6 mice were co-housed with age- and sex-matched CD-1 donors at a 1:1 ratio. Numbers obtained from the co-housing procedure were as follows for the first experiment: GM^High^CH (*n* = 12/sex), GM^Low^CH (*n* = 12/sex). For the second experiment: GM^High^CH (*n* = 7 males, 8 females), GM^Low^CH (*n* = 8 males, 8 females).

#### Gastric gavage (GA)

B6J and B6N mice, obtained directly from respective producers, were bred to generate GM recipient mice. At weaning, mice were placed into cages with littermates of the same sex. Following weaning, mice were exposed to the reciprocal GM by gastric gavage of 0.2 mL of fecal slurry once per week prepared from feces of age- and sex-matched CD-1 donor mice, and transfer of dirty bedding from cages of age- and sex-matched CD-1 donor mice three times per week up to seven weeks of age (**Sup Fig S1I,J**). The fecal slurry was prepared by collecting a fecal sample from respective CD-1 donor mice. For each sample, 1 mL of 1X PBS was added to a 2 mL microcentrifuge tube containing a 5 mm steel ball and an approximately 5 mm portion of the donor fecal pellet. The fecal sample was homogenized in a Qiagen Tissuelyser 2.0 for 30 seconds at 30 Hz to create a fecal slurry. Following homogenization, all samples were pooled within their respective GM by passing the individual slurry samples through a 70 µm nylon mesh filter into the same 50 mL conical tube. Numbers obtained for the gastric gavage groups in the second experiment were as follows: GM^High^GA (*n* = 6 males, 7 females), GM^Low^GA (*n* = 7 males, 6 females).

### Fecal sample collection

Antemortem fecal samples were collected by placing mice in a sterile autoclaved empty cage and allowing them to defecate 2-3 fecal pellets which were promptly collected and stored at −80°C until DNA extraction was performed. Feces were collected from pregnant GM donor CD-1 mice on day 18 of gestation to limit the incidence of cannibalism of the pups. GM recipient mouse fecal samples were collected at three and seven weeks of age.

### Dextran sodium sulfate administration

At seven weeks of age, all recipient mice were administered freshly reconstituted dextran sodium sulfate (DSS) at a concentration of 2.5% in drinking water for seven days, followed by seven days of DSS-free standard autoclaved drinking water. Mice were weighed daily during the seven days of DSS administration and the following seven days after discontinuing DSS to monitor weight loss, with exception of the GM^High^CF cohort where the first- and third-day’s weights during DSS administration were inadvertently not recorded. At the end of the 14 days, mice were humanely euthanized, and samples collected. Per the IACUC protocol humane endpoints, any mice that lost greater than or equal to 20% of their pre-DSS administration weight, or were assessed by the investigators to be moribund, were humanely euthanized and samples immediately collected.

### Necropsy

At nine weeks of age, all DSS-treated mice were humanely euthanized by CO_2_ asphyxiation, followed by cervical dislocation according to the AVMA guidelines on humane euthanasia. Immediately following euthanasia, the cecum and colon were removed, and colon lengths were measured from the cecocolic junction to the rectum. The most distal fecal pellets within the colon were collected and promptly stored at −80°C. For the first experiment, cecum and colons were flushed, placed into cassettes, and immersed in 10% neutral buffered formalin to fix for histological slide preparation. For the second experiment, the colon was incised longitudinally and flattened on card stock, serosal side down. Tissue was then bisected longitudinally and one half of the full length was immersion fixed in formalin while the other half was placed in a 2 mL microcentrifuge tube, flash frozen in liquid nitrogen, and promptly stored at −80°C until protein extraction was performed for immune mediator analysis.

### Microbiome analysis

#### DNA extraction

DNA from fecal samples was extracted using the QIAamp PowerFecal DNA kit (Qiagen) per manufacturer instructions, with the exception that homogenization was performed in a 2 mL microcentrifuge tube containing a 5 mm steel ball and placed in a Qiagen Tissuelyser 2.0 at 30 Hz for 10 minutes. All other steps were performed per the manufacturer instructions. DNA concentration was quantified using the Qubit® 2.0 Fluorometer with the Qubit dsDNA BR assay (Invitrogen) following manufacturer’s protocol.

#### 16S rRNA amplicon library preparation and sequencing

Extracted fecal DNA was processed at the University of Missouri Genomics Technology Core Facility. Bacterial 16S rRNA amplicons were constructed via amplification of the V4 region of the 16S rRNA gene using previously developed universal primers (U515F/806R), flanked by Illumina standard adapter sequences^26,27^. Oligonucleotide sequences are available at proBase^28^. Dual-indexed forward and reverse primers were used in all reactions. PCR was performed in 50 µL reactions containing 100 ng metagenomic DNA, primers (0.2 µM each), dNTPs (200 µM each), and Phusion high-fidelity DNA polymerase (1U, Thermo Fisher). Amplification parameters were 98°C^(3^ ^min)^ + [98°C^(15^ ^sec)^ + 50°C^(30^ ^sec)^ + 72°C^(30^ ^sec)^] × 25 cycles + 72°C^(7^ ^min)^. Amplicon pools (5 µL/reaction) were combined, thoroughly mixed, and then purified by addition of Axygen Axyprep MagPCR clean-up beads to an equal volume of 50 µL of amplicons and incubated for 15 minutes at room temperature. Products were washed multiple times with 80% ethanol and the dried pellet was resuspended in 32.5 µL EB buffer (Qiagen), incubated for two minutes at room temperature, and then placed on a magnetic stand for five minutes. The final amplicon pool was evaluated using the Advanced Analytical Fragment Analyzer automated electrophoresis system, quantified using quant-iT HS dsDNA reagent kits, and diluted according to Illumina’s standard protocol for sequencing on the MiSeq instrument.

#### Bioinformatics

Primers were designed to match the 5’ ends of the forward and reverse reads. Cutadapt^29^ (version 2.6) was used to remove the primer from the 5’ end of the forward read. If found, the reverse complement of the primer to the reverse read was then removed from the forward read as were all bases downstream. Thus, a forward read could be trimmed at both ends if the insert was shorter than the amplicon length. The same approach was used on the reverse read, but with the primers in the opposite roles. Read pairs were rejected if one read or the other did not match a 5’ primer, and an error-rate of 0.1 was allowed. Two passes were made over each read to ensure removal of the second primer. A minimal overlap of three bp with the 3’ end of the primer sequence was required for removal. The QIIME2^30^ DADA2^31^ plugin (version 1.10.0) was used to denoise, de-replicate, and count ASVs (amplicon sequence variants), incorporating the following parameters: 1) forward and reverse reads were truncated to 150 bases, 2) forward and reverse reads with number of expected errors higher than 2.0 were discarded, and 3) Chimeras were detected using the “consensus” method and removed.

#### Tissue histological examination and scoring

Cecum and colon were trimmed, embedded, and sectioned by the histology services of IDEXX BioAnalytics (Columbia, MO). Histological examination was performed by two blinded laboratory animal veterinarians experienced in reviewing GI tissues (KG and TR). Slides were randomly ordered so that reviewers were blinded to transfer method, transfer direction, and sex. Reviewers assigned a lesion score based on the degree of inflammation and epithelial changes, and overall percentage of the colonic lesions (**Sup Table S1**). Scores that differed by 1 between reviewers were averaged. When scores differed by greater than 1, reviewers re-examined slides together and generated a consensus score. Only after agreement was reached on scores were reviewers unblinded to treatment groups.

### Protein extraction and immune mediator analysis

Colon tissue from six males and six females from each GM transfer group in the co-housing and gavage groups were randomly selected for cytokine analysis. Protein was extracted from colon tissue by adding 500 µL of 1X phosphate-buffered saline to each colon tissue sample. Samples were then homogenized in a 2 mL microcentrifuge tube containing a 5 mm-diameter steel ball. Mechanical homogenization was performed using a Qiagen Tissuelyser 2.0 at a frequency of 30 Hz for 5 minutes. Samples were then centrifuged at 9,000 × *g* for 9 minutes and supernatant collected. Protein concentrations were quantified using the Qubit® 2.0 Fluorometer with the Qubit protein BR assay (Invitrogen) following the manufacturer instructions. Protein samples were analyzed at a concentration of 300-550 ug/mL using a ProcartaPlex^TM^ Mouse Immune Monitoring Panel 48-Plex kit (Invitrogen) according to manufacturer instructions. A standard curve was generated using the standards provided and according to the manufacturer’s protocol. All samples were run in duplicate. Data was acquired on a routinely validated and calibrated Luminex xMAP INTELLIFLEX system. Samples for which the immune mediator concentrations were too low to be detected were designated to have a concentration of zero due to the high sensitivity and specificity of the assay.

### Statistics

Two-way permutational analysis of variance (PERMANOVA) was used to test for significant main effects in beta diversity of transfer method and donor/recipient. Three-way PERMANOVA was used to test for significant main effects in beta diversity of transfer method, transfer direction, and donor/recipient, followed by one-way PERMANOVA for donor/recipient group pairwise comparisons. One-way and Two-way PERMANOVA analysis was performed using PAST 4.09 software^32^ and was based on Jaccard dissimilarities. Three-way PERMANOVA testing was based on Jaccard dissimilarities using the *adonis2* library from the *vegan* library^33^. Distances between group centroids were determined using the *usedist* library^34^. Comparisons in percent weight change were performed by calculating area under the curve (AUC) for each mouse from days 6 to 14 of DSS treatment when marked weight loss occurred, and normalizing AUC to days survived to account for animals that were removed from the study during DSS treatment due to reaching humane endpoints. For percent weight loss data in experiment 1, a one-way analysis of variance (ANOVA) was used to test for effect of transfer method within the GM^High^ cohorts followed by Tukey *post hoc* for pairwise comparisons. Due to lack of normality in the GM^Low^ cohorts, a Kruskal-Wallis ANOVA on Ranks was used to test for effect of transfer method followed by Dunn’s *post hoc* for pairwise comparisons. A Student’s t-test was used to test for significant differences between the percent weight loss of GM^High^CH and GM^Low^CH. Due to lack of sufficient animals remaining in the GM^Low^CH cohort (two animals remained following day 10), statistical analysis of the cohort percent weight loss was only performed to day 10 of DSS treatment. For percent weight loss data in experiment 2, a two-way ANOVA was used to test for main effects of transfer method and transfer direction followed by Tukey *post hoc* for pairwise comparisons. For unifactorial survival data, a survival LogRank analysis was used to test for significant effects. For multifactorial survival data, a Cox proportional hazards test was used to test for significant main effects in disease survivability including transfer method, transfer direction, and sex. Three-way ANOVA was used to test for significant main effects of transfer method, transfer direction, and sex for Chao-1 richness, weaning and week seven weights, colon lengths, histological lesion scores, and cytokine/chemokine concentrations followed by Tukey’s *post hoc* analysis for pairwise comparisons. Univariate data was first tested for normality using the Shapiro-Wilk method. All univariate data analysis was performed using SigmaPlot 15.0 (Systat Software, Inc, San Jose, CA). Due to lack of normality in the concentrations of MIP-2α, IL-22, and IL-6 these data were transformed prior to performing three-way ANOVA analysis. MIP-2α concentrations were normalized by square root transformation, and IL-22 and IL-6 concentrations were normalized by logarithmic (log) transformation. Due to uniform lack of normality across immune mediator concentrations, a Mann-Whitney U test was used to test for statistical differences in immune mediator concentrations between GM^High^ and GM^Low^ treatment groups, as well as co-housing and gavage treatment groups. All diversity and richness indices were calculated using PAST 4.09 software.

## Results

### Efficiency of GM transfer is determined by transfer method

We first sought to determine if we could replicate our previous findings^22^ in an acute DSS-colitis disease model. To provide clarity regarding transfer terminology and nomenclature used in this study, we have provided **Sup Fig S1** as a schematic to assist the reader in following the experimental groups and transfer procedures. To be clear, all GM donor CD-1 mice used in this study were from outbred colonies where the founders were originally purchased from Charles River, and given their respective microbiomes via ET^24^. To assess GM transfer efficiency, transfer recipients receiving the respective GM via ET were compared to the CD-1 ET dams (i.e., the GM donors in this case). Similarly, CF recipients were compared directly to their CF surrogate dams. For CH, recipients were compared to feces from the dams of the CD-1 co-housing donors as the GMs of the CD-1 donors housed with the B6 recipients would be modulated by the recipient B6 microbiome via mutual coprophagia. We performed 16S rRNA amplicon sequencing analysis on feces from the recipient mice of the ET, CF, and CH groups at seven weeks of age, and compared these to the respective donor fecal microbiome. To assess transfer efficiency, unweighted beta diversity between donors and recipients was first examined. Principal coordinate analysis (PCoA) revealed little separation between recipient and donors in the GM^High^ET, CF, and CH groups, with the GM^High^CH group showing greater dissimilarity to their donors (Distance between centroids [CD] = 0.500) compared with the ET (CD = 0.409) and CF (CD = 0.283) groups (**Figure 1a**). In contrast, the GM^Low^ET and CF groups showed similar composition to their donors (CD of 0.322 and 0.386, respectively), but the GM^Low^CH group showed marked separation from their donors (CD = 0.623) (**Figure 1b**). When beta diversity of all six transfer groups was compared, a stark difference in beta diversity between the recipient and donors within the GM^Low^CH group and the other transfer groups became apparent by the greater separation of the GM^Low^CH donors and recipients (**Sup Fig S2; Sup Table S3**). When alpha-diversity was assessed, the GM^High^ recipients showed no statistically significant differences in GM richness between the donor and recipient GMs in any transfer method (**Figure 1c**). The GM^Low^ET and CF groups showed no significant differences between donor and recipient mice, but a marked significant difference was observed in the GM^Low^CH group with the recipient mice having a greater GM microbial richness compared with the donors at seven weeks of age (**Figure 1d**) resulting in a pattern similar to what was observed with beta diversity.

**Figure 1.**
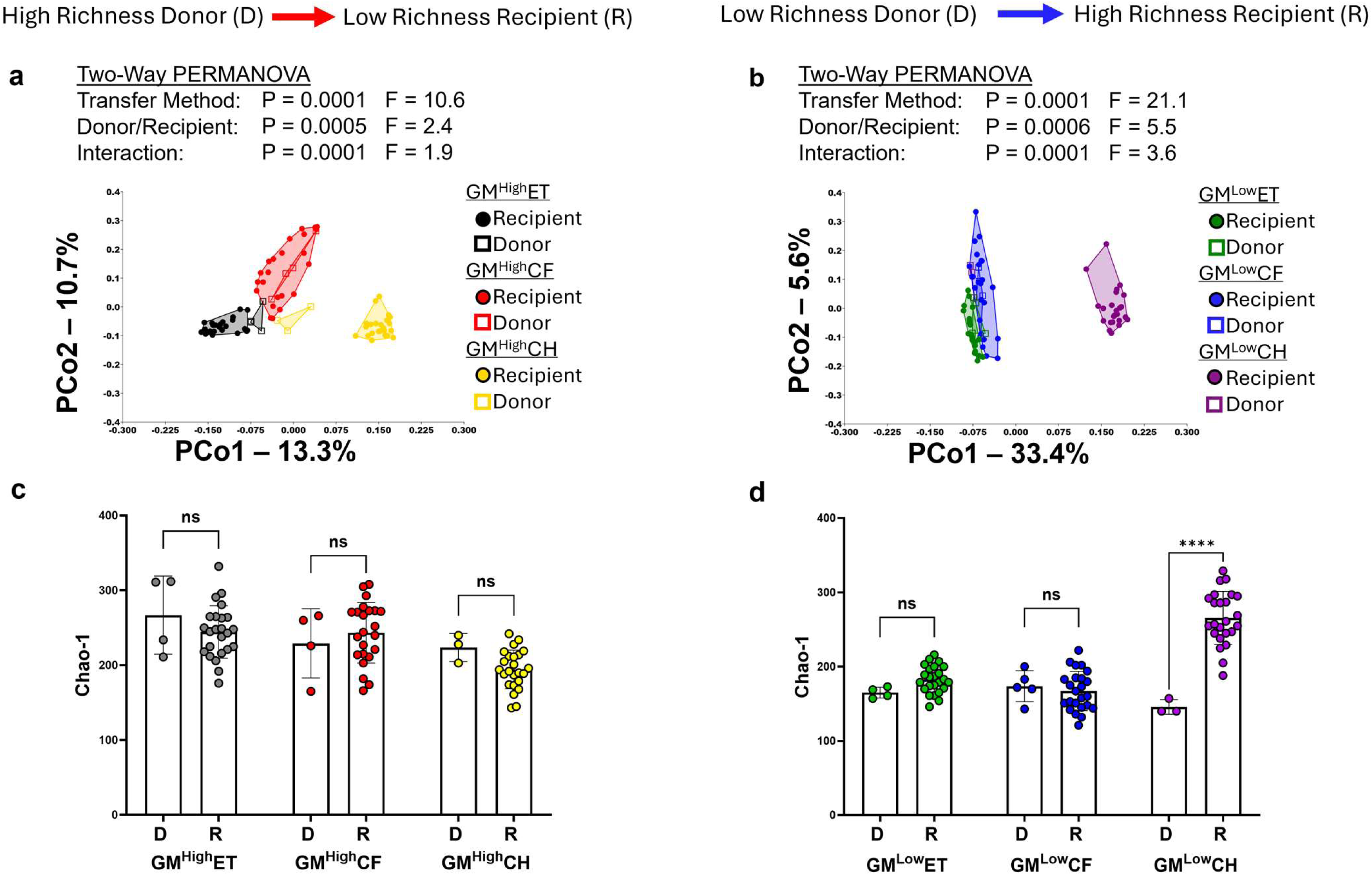
Characterization and comparison of the six transfer groups to determine efficiency of GM transfer. Principal coordinate analysis comparing the three transfer methods of the (**a**) GM^High^ and the (**b**) GM^Low^ group’s beta diversity to their donors. X and Y axes labeled with percent of variation contributed by Principal coordinate 1 (PCo1) and PCo2, respectively. Two-way PERMANOVA for main effects of transfer method and recipient/donor (**Sup Table S2**), followed by a one-way PERMANOVA to test pairwise comparisons between donor and recipient (**Sup Table S3**). Comparison of donor and recipient GM at seven weeks of age for the (**c**) GM^High^ and (**d**) GM^Low^ cohorts. Bars represent mean chao-1 richness +/- SD. Two-way ANOVA for main effects of transfer method and sex. **ns** - not significant, **** p<0.0001.

### Disease phenotype determined by efficiency of GM transfer

#### Weight loss

While the GM^High^ groups showed a small but significant difference in weight loss between the CF and CH groups (**Figure 2a**), the GM^Low^ cohorts showed a dramatic significant difference in their weight loss between the three groups with CH have the most severe weight loss (**Figure 2b**). Of note, while pre-DSS body weights collected at weaning revealed significant differences between transfer method groups in both GM^High^ and GM^Low^ recipient mice (**Sup Figure S3A-B**), those differences were normalized at seven weeks of age, the age when DSS administration began, with no differences detected between groups (**Sup Fig 3C-D**). While CH was clearly associated with more severe disease in GM^Low^CH mice, GM^High^CH mice developed similar disease to GM^High^ET and GM^High^CF mice, suggesting that CH *per se* is not solely responsible for the severe disease observed in GM^Low^CH mice (**Sup Fig S4A-B**).

**Figure 2.**
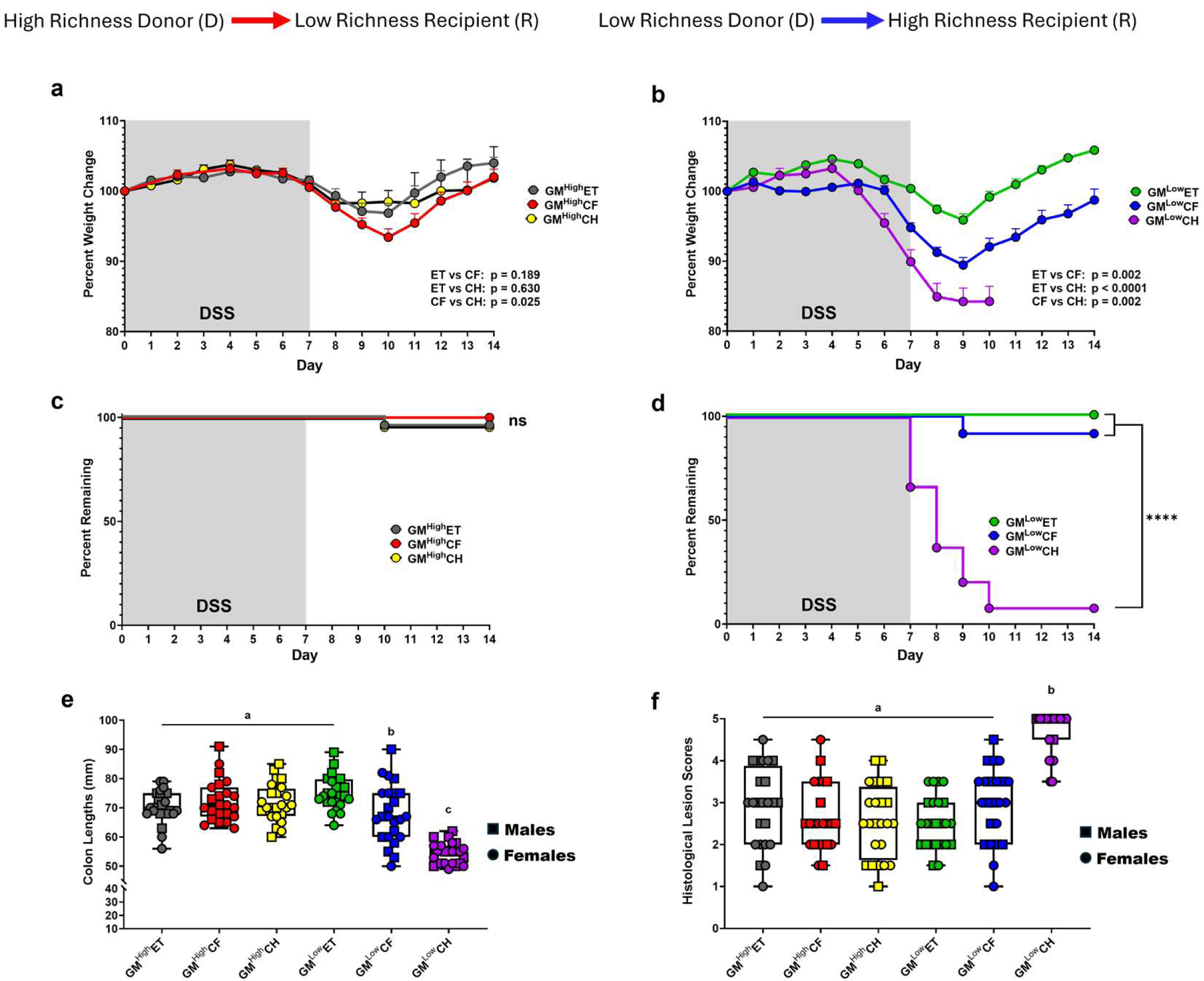
Comparison DSS-colitis disease phenotype differences of the six transfer groups. (**a**) Comparison of the DSS-induced weight loss between the GM^High^ cohort transfer methods. One-way ANOVA for effect of transfer method (p = 0.013, F = 4.5). Tukey *post hoc* for pairwise comparisons. (**b**) Comparison of the DSS-induced weight loss between the GM^Low^ cohort transfer methods. Kruskal-Wallis ANOVA on ranks for effect of transfer method (p <0.0001, F = 51.4). Dunn’s *post hoc* for pairwise comparisons. Each data point in panels (**a**) and (**b**) represents transfer method mean percent weight change +/- SEM. DSS-induced disease survivability of the (**c**) GM^High^ cohorts and the (**d**) GM^Low^ cohorts. Cox proportional hazards for main effects of transfer method and sex. (**e**) Colon lengths for each cohort following DSS administration and (**f**) DSS-induced lesion scores for each cohort. Groups that differ in letter designation are statistically significant from each other, while groups that share the same letter designation are not statistically significant from each other. Three Way ANOVA for main effects of transfer method, transfer direction, and sex. **ns** - not signification, **** p<0.0001.

#### Survival

To assess whether the method and efficiency of GM transfer can influence disease phenotype, a DSS colitis model was employed in which DSS was administered in the drinking water for one week, followed by one week of recovery with untreated water. Immediate outcomes measures including weight loss and survival demonstrated profound differences in co-housed groups. No significant differences in experimental survival among the GM^High^ groups were observed with only two individuals, one in the ET group and one in the CH group, requiring euthanasia due to disease severity (**Figure 2c**). However, the GM^Low^CH group had a significant number of individuals (91.6%) requiring euthanasia due to weight loss and disease severity (**Figure 2d**).

#### Colon lengths and lesion scores

Following euthanasia, colon length measurements revealed a difference between the GM^Low^CH group, GM^Low^CF groups, and the four other groups (**Figure 2e**) with GM^Low^ CH having the greatest reduction in colon length. To determine if the shorter colon lengths in GM^Low^CH mice were due to being euthanized before the two-week endpoint, colons of mice taken down early were compared to those who continued to the end of study. Interestingly, no difference was detected in colon length of those mice that were euthanized early compared with those of the mice who survived to the end of study (**Sup Fig S5A**). Histological examination also showed the GM^Low^CH group had significantly greater disease with a majority of the lesions characterized by severe colonic epithelial ulceration and erosion with marked immune cell infiltration (**Figure 2f**), the most severe being in those mice that were euthanized early (**Sup Fig S5B**).

### Co-housing disease phenotype is a result of the transferred GM and transfer efficiency

With the interesting repeatable results obtained in the GM^Low^CH group, we next wanted to confirm that these observations were a result of the GM-associated microbial factors and not other factors associated with CH. To this end, a second experiment was performed where GM^High^CH and GM^Low^CH were compared to two groups in which the recipient mice were exposed to reciprocal GMs via gastric gavage (GA) and dirty bedding transfer. Following four weeks of GM transfer via CH or GA, sequencing of the fecal DNA showed that the effect of transfer direction was significant, but not method of transfer, as no differences in microbial composition or richness were found between the CH and GA treatment groups in either transfer direction (**Figure 3a-b**). Similar to previous experiments, transfer of a high richness GM to a recipient harboring a low richness GM resulted in more efficient colonization than transfer of a low richness GM to a recipient harboring a high richness GM (**Sup Fig S6A-C**). When transferring a high richness GM to a low richness recipient by CH or GA, the recipients shared similar GM composition across these transfer methods (**Sup Fig S6A**), and while richness was significantly different, the CH and GA groups had a greater microbial richness than the donors at week seven (**Sup Fig S6B**). In contrast, GM^Low^ donors did not successfully transfer their GM composition or taxonomic richness to the B6N recipients (**Sup Fig S6A,C**).

**Figure 3.**
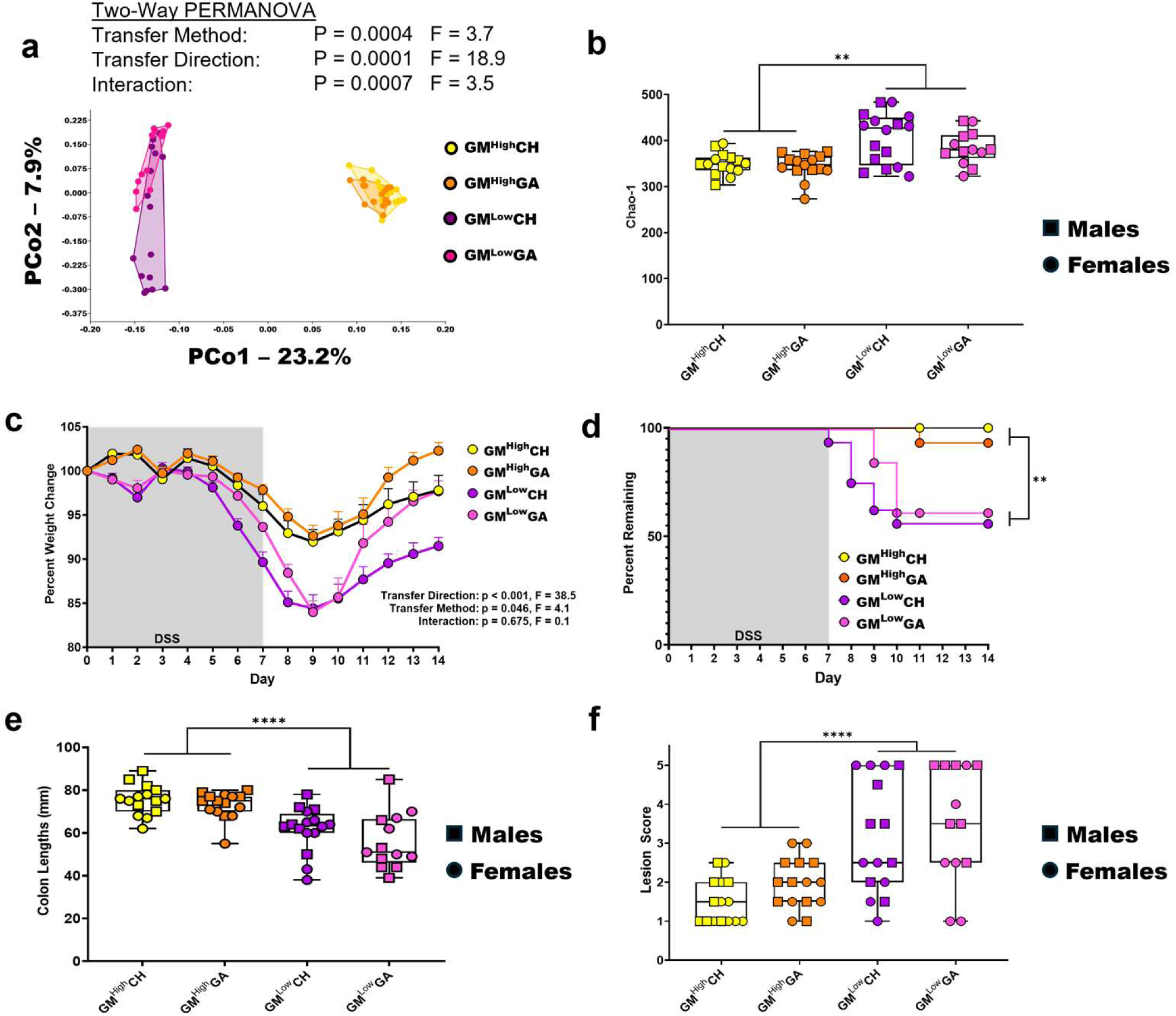
Comparison of co-housing and gavage GM transfer efficiency and DSS-induced DSS phenotype in each transfer direction. (**a**) PCoA comparing the GM beta diversity of the CH and GA cohorts in each transfer direction. X and Y axes labeled with percent of variation contributed by PCo1 and PCo2, respectively. Two-way PERMANOVA for main effects of transfer method and transfer direction. (**b**) Chao-1 richness of each co-housing and gavage cohorts. Three-way ANOVA for main effects of transfer method, transfer direction, and sex. (**c**) DSS-induced weight loss comparison of the co-housing and gavage cohorts. Each point represents group mean percent weight change +/- SEM. Two-way ANOVA for main effects of transfer method and transfer direction. (**d**) DSS-induced disease survivability of the co-housing and gavage cohorts. Cox proportional hazards for main effects of transfer method, transfer direction, and sex. (**e**) Colon lengths and (**f**) histological lesion scores caused by DSS-induced colitis. Three-way ANOVA for main effects of transfer method, transfer direction, and sex. **p<0.01, ***p<0.001, ****p<0.0001.

We next administered DSS to the four treatment groups with a single one-week pulse followed by a week of recovery. A significant difference was observed between the mice receiving GM^Low^ or GM^High^, with transfer of a low richness GM to recipients harboring GM^High^ resulting in greater weight loss (**Figure 3c**) and higher mortality (**Figure 3d**). However, these marked weight loss and survival differences were not observed between the CH and GA groups within either transfer direction. Similarly, analysis of colon lengths collected at necropsy showed a significant difference between GM^Low^ and GM^High^ recipients, while transfer method had no effect within either GM (**Figure 3e**). Colon tissue was also collected for histological examination, which revealed a similar pattern where mice receiving GM^Low^ had significantly greater lesion severity than mice receiving GM^High^, regardless of transfer method (**Figure 3f**).

### Transfer of GM^Low^ to mice harboring GM^High^ results in increased immune mediators within diseased colons

Lastly, local immune responses were assessed using bead-based immunoassay quantification of 45 cytokines, chemokines, and other immune mediators. When concentrations were compared based on the GM being transferred, nine inflammatory mediators were significantly elevated in mice receiving GM^Low^ (**Figure 4a; Sup Table S4**). Alternatively, comparison based on transfer method failed to detect a difference in any of the immune mediator concentrations (**Figure 4b; Sup Table S5**), suggesting that the transfer efficiency, not transfer method, is driving the immune response. Affected cytokines and chemokines included the chemokine MIP-2α (**Figure 4c**) involved in recruitment of innate immune cells in acute phase immune responses, IL-22 (**Figure 4d**) which promotes epithelial proliferation and regeneration in response to injury, and the pro-inflammatory cytokine IL-6 (**Figure 4e**). Immune mediator group concentration means and standard deviations are provided in **Sup Table S6**.

**Figure 4.**
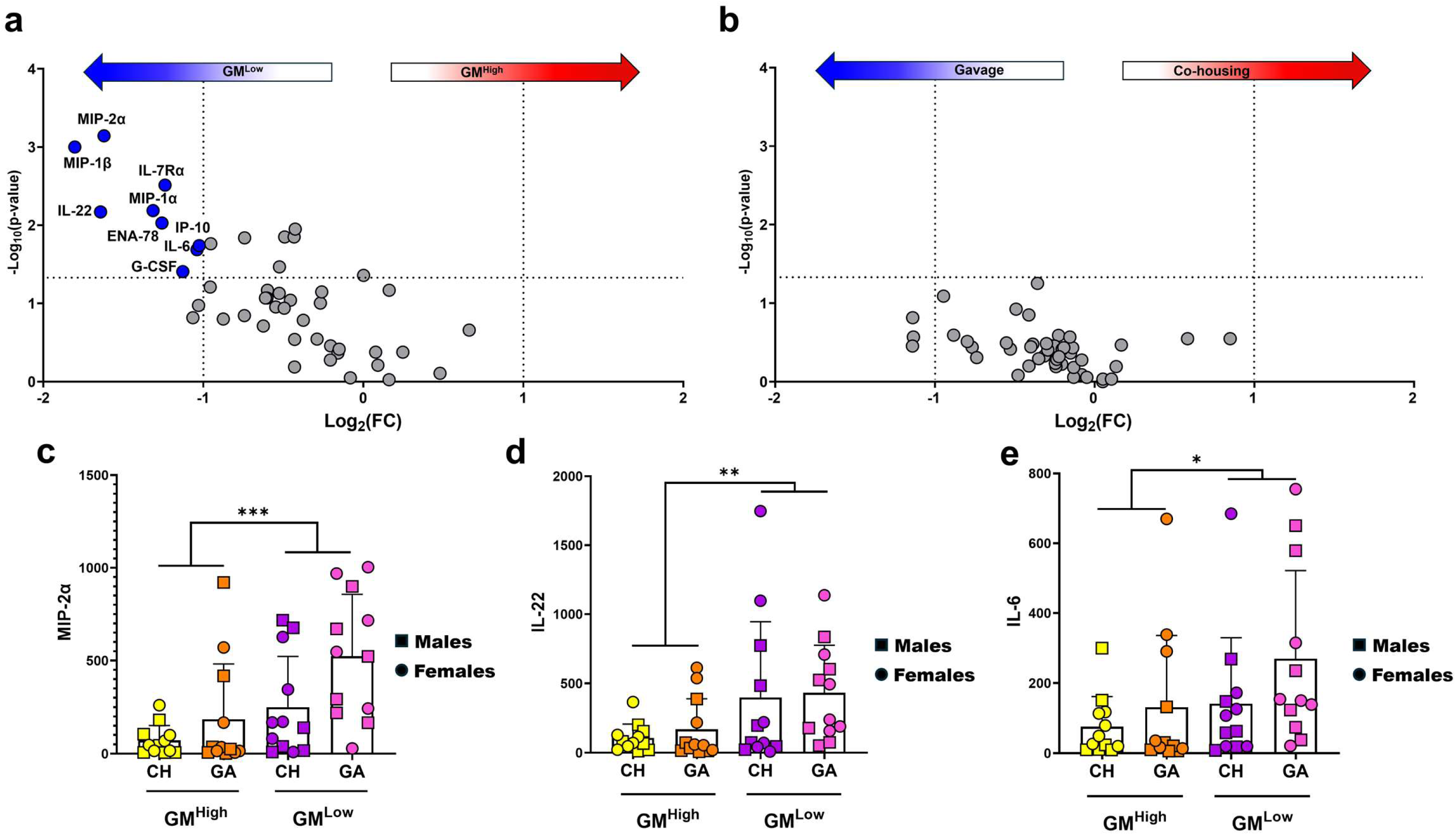
Immune mediator concentrations from the co-housing and gavage cohorts’ colons following DSS-induced colitis. Volcano plots comparing the cytokine and chemokine concentrations between (**a**) GM^High^ and GM^Low^ cohorts, and (**b**) gavage and co-housing cohorts. Immune mediator concentration comparison of the co-housing and gavage cohorts for (**c**) macrophage inflammatory protein 2-alpha (MIP-2α), (**d**) interlukin 22 (IL-22), and (**e**) interlukin 6 (IL-6). Bars represent mean immune mediator concentrations +/- SD. Three-way ANOVA for main effects of transfer method, transfer direction, and sex. *p<0.05, **p<0.01, ***p<0.001.

## Discussion

In this study, we have leveraged an acute model of DSS colitis to elucidate the impact of gut microbiota (GM) composition, richness and efficiency of transfer methods on disease phenotypes. The results presented here confirm previous results that embryo transfer and cross-fostering share similarly high transfer efficiency regardless of GM composition^22^. In contrast, co-housing at weaning is less effective, suggesting that GM transfer early in life facilitates microbial colonization. Furthermore, the relationship of the recipient and donor microbiome during co-housing determines transfer efficiency. Differences between groups in disease severity suggest that the efficiency of transfer influences the severity of weight loss, mortality, reduction in colon length, and histological lesion severity, corroborating previous results using a chronic DSS colitis model^22^. The addition of groups receiving GM transfer via gavage and dirty bedding provides further evidence that the severity of colitis is negatively associated with the efficacy of GM colonization following transfer.

During the colonic epithelial barrier disruption induced by DSS, the presence of non-colonizing microbes is associated with a more robust immune response than bacteria against which the host has presumably been tolerized due to successful colonization and immune recognition. Immune tolerance develops early in life when the host is first introduced to microbes. This first introduction allows the immune system to develop a tolerance through development of Th2 response and expansion of RORgt^+^ T helper lymphocytes and dendritic cells within the gut early in life^35^. These RORgt^+^ cell populations quickly decline soon after birth^36^, and may explain the mild disease severity in the ET and CF groups regardless of GM composition. The relatively mild disease observed in the GM^High^CH and GM^High^GA groups suggests that tolerance is induced in adolescent recipient mice, assuming there is successful colonization of the transferred GM.

Collectively, these findings suggest that administration of DSS and subsequent induction of epithelial barrier defects can be used experimentally to assess colonization and immune recognition of microbial exposures. Molecular methods may show evidence of microbial exposures in the fecal DNA regardless of patent colonization. DSS-induced colitis provides a disease phenotype driven by the immune response to the GM, including prior recognition of antigens within the gut lumen.

The use of FMT to treat diseases including Crohn’s disease, ulcerative colitis, and *Clostridioides difficile* infection has been widely reported^37–39^. The use of a stool sample from a healthy donor to restore a dysbiotic gut microbial community has shown great promise as a treatment. However, there are instances in which FMT has been ineffective or even exacerbated IBD in patients^40–42^. Similarly, probiotics may be used to treat or prevent dysbiosis in IBD patients. The response to probiotics can also be variable and clinical trials show that probiotics can exacerbate IBD symptoms^43^. The present data may provide a partial explanation for these adverse outcomes following FMT or probiotic administration in IBD patients. In the context of disease-associated epithelial damage, recognition of foreign antigens against which the host has not developed tolerance may exacerbate disease.

### Our study is not without its limitations

For the first experiment in this study, it was not possible to truly determine the extent to which the co-housing donors transferred their microbiomes, as weaning samples from the donors do not represent a fully developed GM of an adult and the seven-week samples were collected after co-housing changed the donor microbiome by transfer of microbes from the recipient mouse GM. However, as it has been shown that the dam will efficiently transfer her GM to her pups soon after birth, the GM harbored by the dam of the CH donor mice will be a sufficient representation of the donor GM. For the immunoassay, we only measured 45 immune mediators, while we recognize that there are many more that can influence disease severity. That said, well-characterized immune mediators were included in the assay, both pro-inflammatory and anti-inflammatory. We also recognize that in performing weekly gastric gavage, we introduced momentary acute stress by handling the mice. However, the goal of the weekly gavage transfer was to eliminate physical contact and effects of social interaction.

### Conclusion

In summary, our findings indicate that colonization efficiency following GM transfer is determined by the relationship between donor and recipient GM, and that poor transfer efficiency is associated with more severe disease. Moreover, these data provide further evidence that methods used to manipulate the GM must be considered in the context of study reproducibility when results between similar studies are not in agreement.

## Acknowledgments

We would like to thank Dr. Rachel Olson, Dr. James Chung, and Amy Steeneck for their knowledge and technical assistance with immune mediator concentration collection and analysis. We would also like to thank Benjamin Olthoff for his technical assistance with DSS preparation and administration, and Rebecca Dorfmeyer for her assistance with fecal DNA sample preparation for 16S rRNA amplicon sequencing analysis.

## Funding Statement

KG, ZM, ACE, and CLF and project supplies and animals were supported by NIH U42 OD010918. KG was also supported by NIH T32 OD011126 and the Joseph Wagner Fellowship Endowment in Laboratory Animal Medicine and ZM was also supported by NIH T32 GM008396. TR was supported by the University of Missouri Office of Research, Innovation and Impact, and the Joseph Wagner Fellowship Endowment in Laboratory Animal Medicine.

## Disclosure statement

The authors report there are no competing interests to declare.

## Data availability statement

All 16S rRNA amplicon sequencing data are available at the NCBI Sequence Read Archive under the BioProject number PRJNA1031529.

## Supplemental Figure Legends

**Sup Fig S1.**
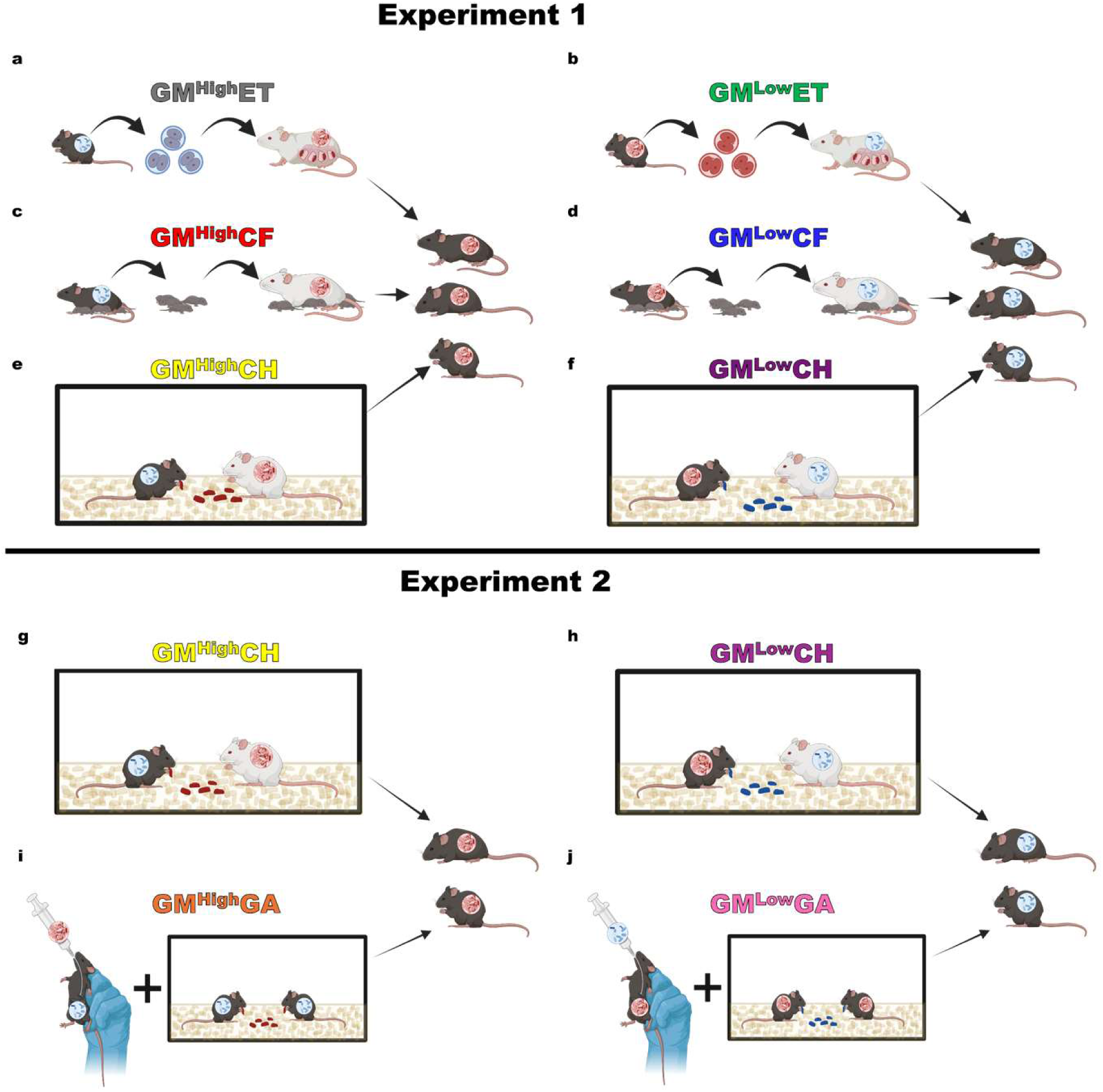
Schematic of the generation of each GM transfer cohort used in the study. (**a**) 2-cell stage embryos were collected from B6J mice and implanted into pseudopregnant CD-1 mice harboring GM^High^. (**b**) 2-cell stage embryos were collected from B6N mice and implanted into pseudopregnant CD-1 mice harboring GM^Low^. (**c**) Pups of less than 24-hours were cross-fostered from B6J dams onto a CD-1 dam harboring GM^High^. (**d**) Pups of less than 24-hours were cross-fostered from B6N dams onto a CD-1 dam harboring GM^Low^. (**e**) B6J mice were weaned into cages with weaned CD-1 donor mice harboring GM^High^ so that the CD-1 donors would transfer their GM^High^ via coprophagia to the B6J mice. (**f)** B6N mice were weaned into cages with weaned CD-1 donors harboring GM^Low^ so that the CD-1 mice would transfer their GM^Low^ via coprophagia to the B6N mice. (**g**) Similar to experiment 1, B6J mice were weaned into cages with weaned CD-1 donor mice harboring GM^High^ so that the CD-1 donors would transfer their GM^High^ via coprophagia to the B6J mice. (**h)** Similar to experiment 1, B6N mice were weaned into cages with weaned CD-1 donors mice harboring GM^Low^ so that the CD-1 donors would transfer their GM^Low^ via coprophagia to the B6N mice. (**i**) B6J mice were gastric gavaged with fecal material from GM^High^ donors beginning at weaning and exposed to GM^High^ via dirty bedding transfer from cages housing the GM^High^ CD-1 GM donors to allow GM transfer via coprophagia. (**j**) B6N mice were gastric gavaged with fecal material from GM^Low^ donors beginning at weaning and exposed to GM^Low^ via dirty bedding transfer from cages housing the GM^Low^ CD-1 GM donors to allow GM transfer via coprophagia. For clarity, the white CD-1 mice in the schematic are the GM donors, and the recipient mice are the black B6 mice to the right of the CD-1 donors.

**Sup Fig S2.**
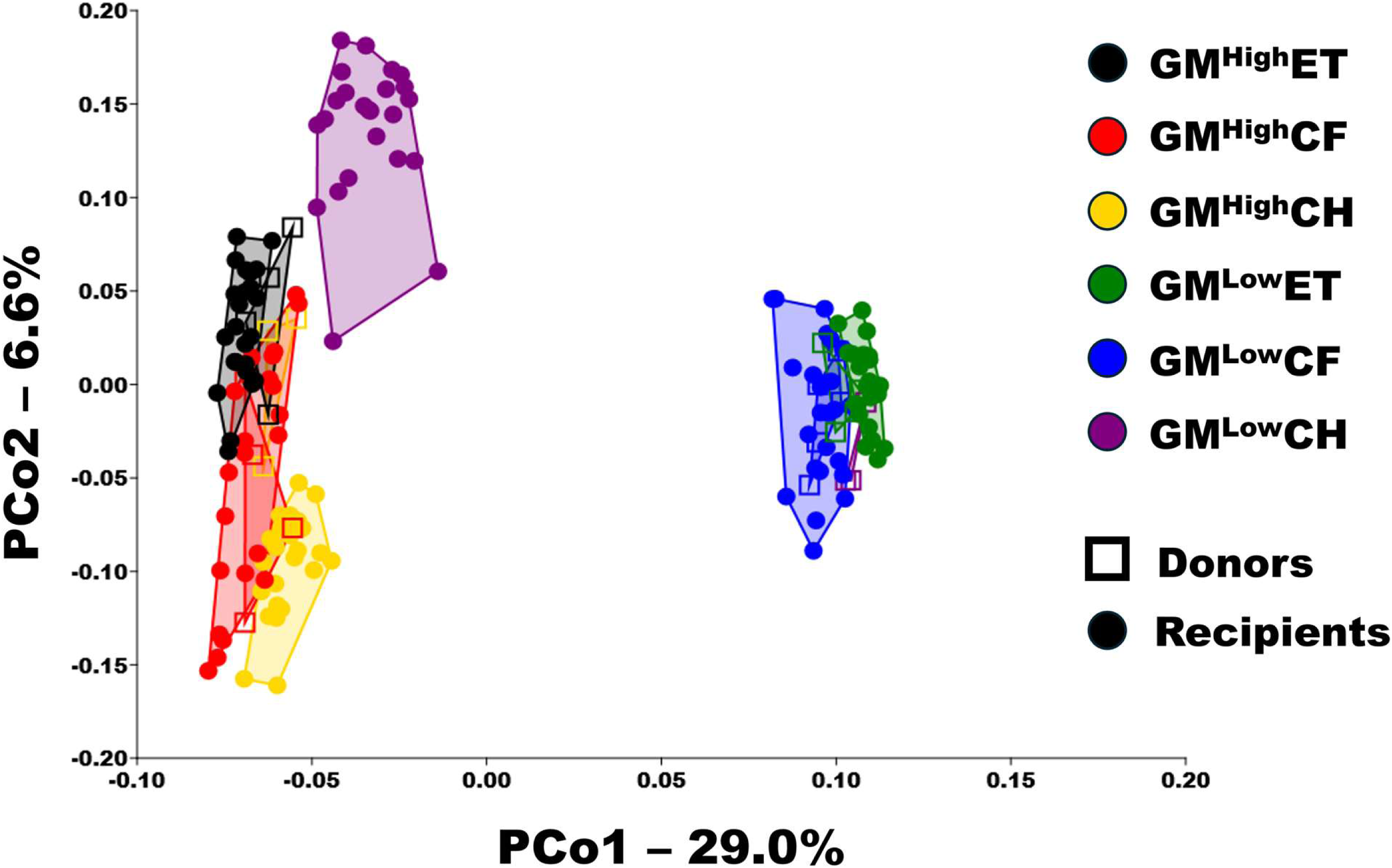
PCoA comparing the six treatment groups’ GM beta diversity to the beta diversity of the donors. X and Y axes labeled with percent of variation contributed by Principal coordinate 1 (PCo1) and PCo2, respectively. Three-way PERMANOVA for main effects of transfer method, transfer direction, and donor/recipient (**Sup Table S2**), followed by a one-way PERMANOVA to test pairwise comparisons between donor and recipient (**Sup Table S3**).

**Sup Fig S3.**
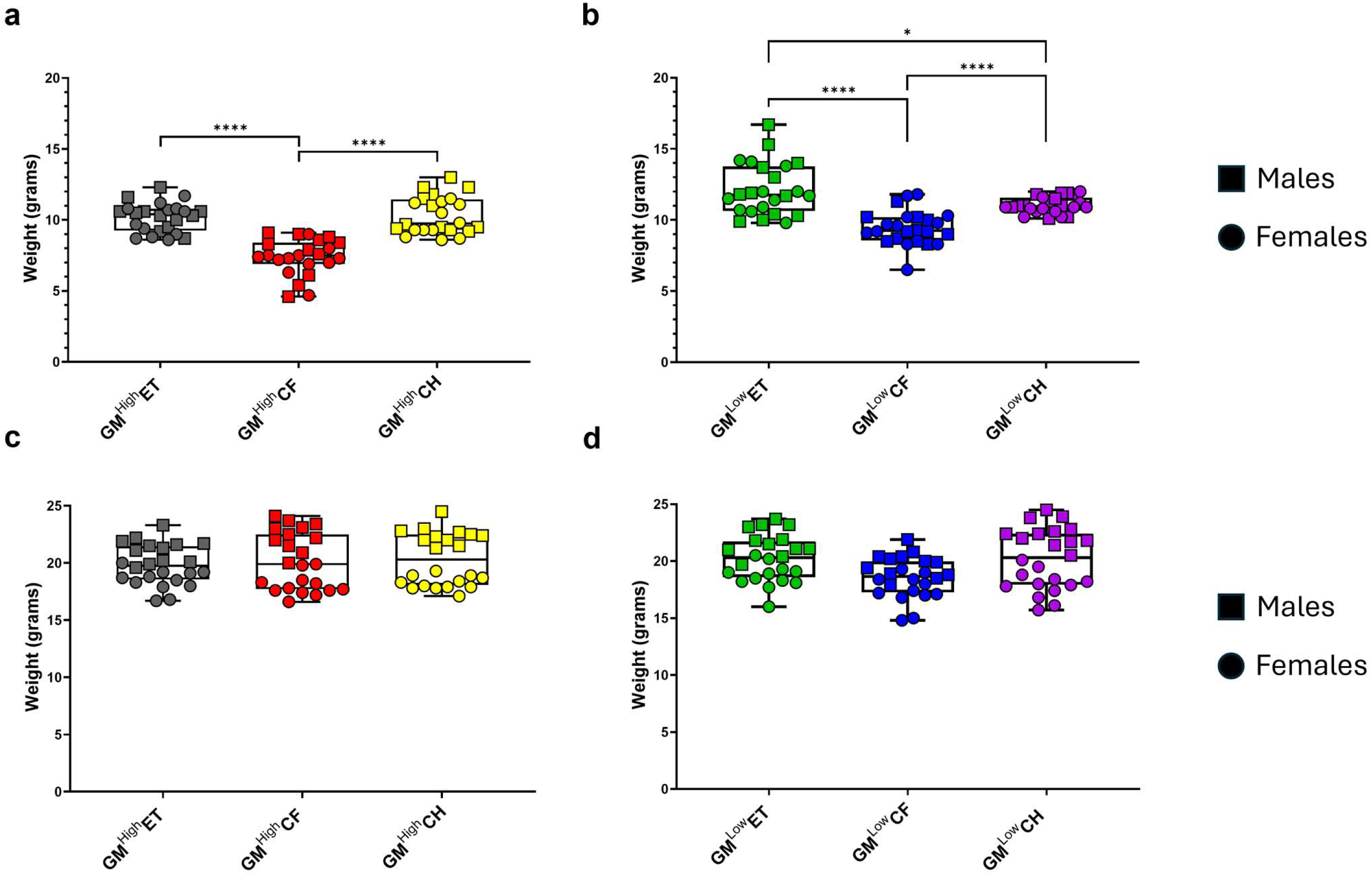
Weaning and week seven weights of the six ET, CF, and CH cohorts. Weaning weights collected from the three (**a**) GM^High^ cohorts and the three (**b**) GM^Low^ cohorts. Weights collected at seven weeks of age prior to DSS administration of the three (**c**) GM^High^ cohorts and the three (**d**) GM^Low^ cohorts. Two-way ANOVA for main effects of transfer direction and sex. *p<0.05, ****p<0.0001.

**Sup Fig S4.**
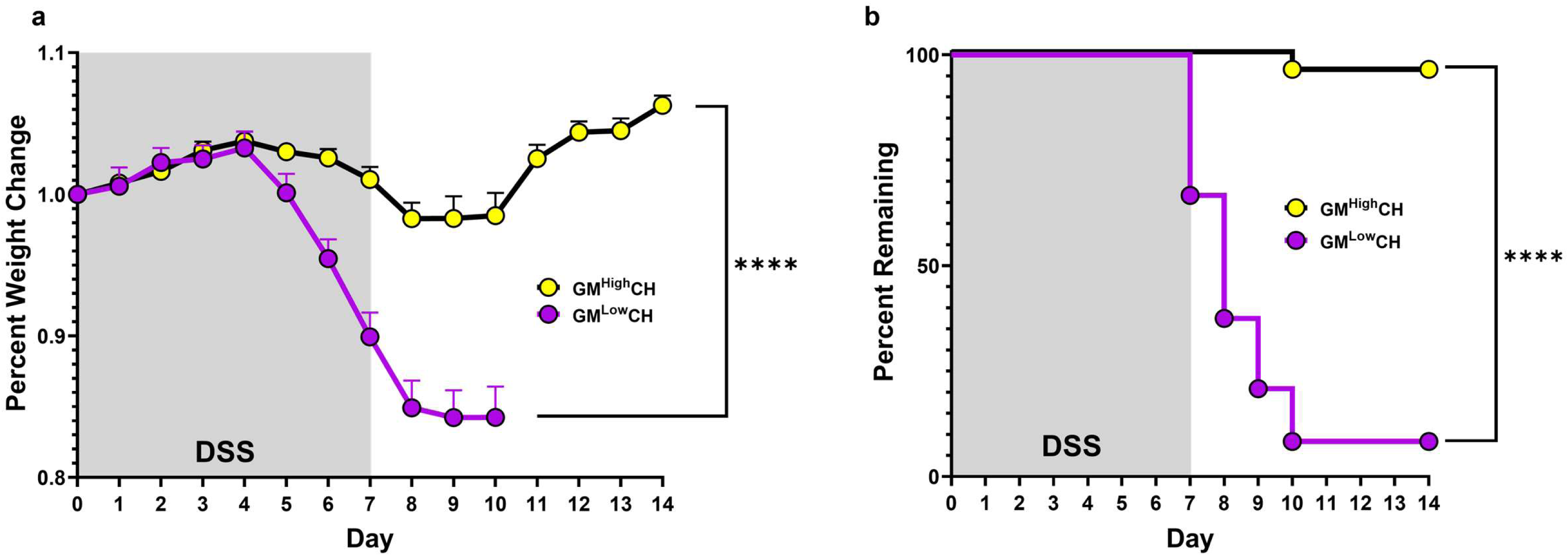
Comparison of the GM^High^CH and GM^Low^CH weight loss and disease survivability phenotypes. (**a**) DSS-induced weight loss comparison of the two co-housing cohorts. Each data point represents group mean percent weight change +/- SEM. Students t-test. (**b**) DSS-induced disease survivability of each co-housing cohort. Survival LogRank. ****p<0.0001.

**Sup Fig S5.**
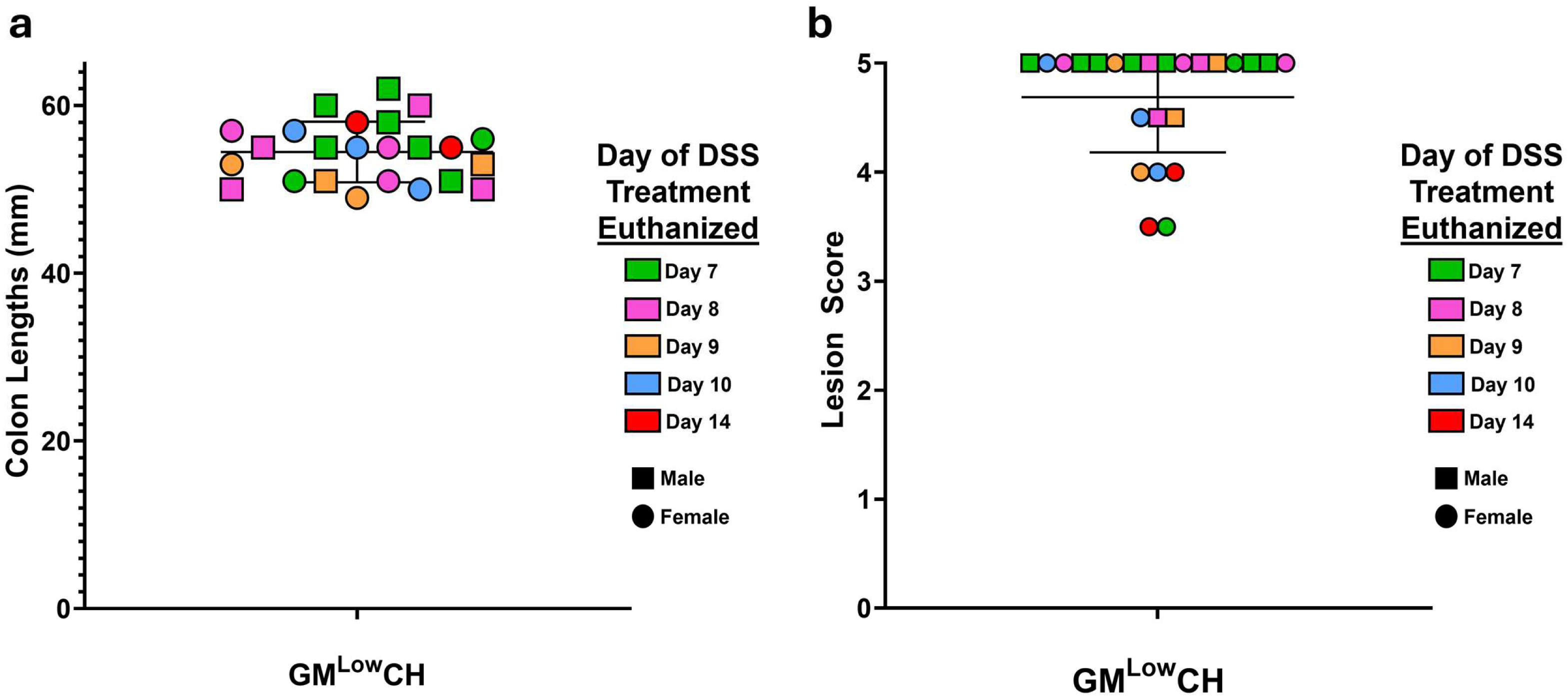
Colon length is not determined by day of DSS-induced colitis treatment at which animals needed to be euthanized, but histological lesion score is dependent on day of DSS treatment in the GM^Low^CH cohort. (**a**) Comparison of the day that animals need to be euthanized due to weight loss (denoted by color), and the length of colons measured at time of necropsy. (**b**) Comparison of the day that animals need to be euthanized due to weight loss (denoted by color), and histological lesion score of the colons.

**Sup Fig S6.**
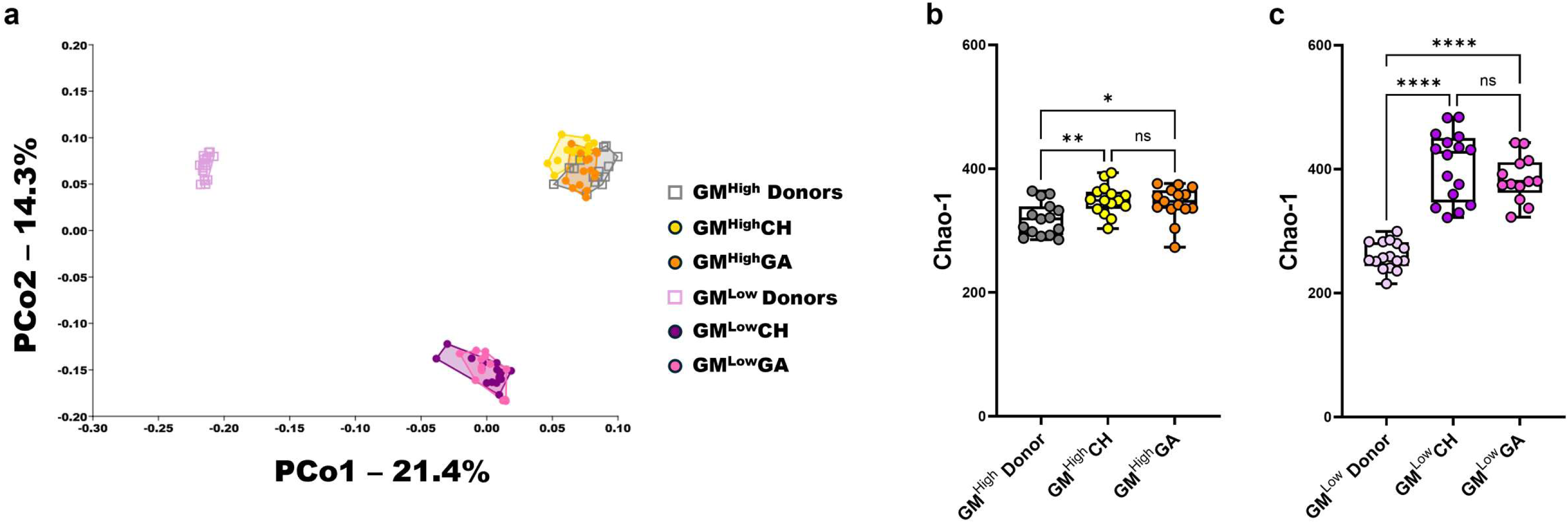
The pattern of transferring the GM at time of weaning leading to a decreased efficiency of transfer is repeatable. (**a**) Principal coordinates analysis comparing the GM^High^ and GM^Low^CH and GA groups to their donor GMs. X and Y axes labeled with percent of variation contributed by PCo1 and PCo2, respectively. Two-way PERMANOVA for main effects of transfer method and group (**Sup Table S7**), followed by a one-way PERMANOVA to test pairwise comparisons between respective donors and recipients (**Sup Table S8**). Chao-1 richness of the (**b**) GM^High^ CH and GA cohorts to the GM^High^ donors, and the (**c**) GM^Low^ CH and GA cohorts to the GM^Low^ donors. Two-way ANOVA for main effects of transfer method and donor/recipient. **ns** - not significant, * p<0.05, ** p<0.01, **** p<0.0001. For the **Sup Fig S6 figures**, we chose to use the GA donors from both transfer directions to represent the GM the recipients should have received (Both CH and GA) as the CH donors’ GM was modified by the recipients’ GM during co-housing and is therefore not a proper representation of the GM alpha- and beta-diversity we wanted the recipients to receive. The GA CD-1 donors were housed separate of their recipients and were not exposed to the GMs of the B6J or B6N mice, and thus were a better representation of what the recipients’ GM should look like if GM transfer was successful. For statistical purposes, the main effect of group in the two-way ANOVA consisted of the donor group compared with the CH and GA recipients pooled as one group within transfer direction. We used a one-way PERMANOVA for individual pairwise comparisons of the beta diversity between the GA donors and CH and GA recipients, and one-way ANOVA to compare richness of the GA donors to the CH and GA recipients.

## Supplemental Table Legends

**Sup Table S1**. Histological lesion scoring criteria used to assign lesion score to DSS treated colons.

**Sup Table S2**. Results of three-way PERMANOVA comparing beta-diversity of ET, CF, and CH groups and the donors, with main effects including transfer method, transfer direction, and donor/recipient.

**Sup Table S3**. Results of one-way PERMANOVA pairwise comparisons of the six ET, CF, and CH recipients and their respective donors. Jaccard dissimilarity distances were measured from recipient group centroid to the centroid of the respective donor group.

**Sup Table S4**. Immune mediatory group averages and results of Mann-Whitney U Test between GM^High^ and GM^Low^ cohorts for immune mediator concentrations.

**Sup Table S5.** Immune mediator group averages and results of Mann-Whitney U Test between co-housing and gavage cohorts for immune mediator concentrations analyzed.

**Sup Table S6.** Cohort means and standard deviations for all immune mediators analyzed.

**Sup Table S7**. Results of the two-way PERMANOVA comparing beta-diversity of CH and GA cohorts to their respective donors, with main effects including transfer method and donor/recipient.

**Sup Table S8**. Results of one-way PERMANOVA pairwise comparisons of the CH and GA cohorts beta diversity with their respective donors.

## Notes

### Competing Interest Statement

The authors have declared no competing interest.

